# A Generalized Discrete Dynamic Model for Human Epidemics

**DOI:** 10.1101/2020.02.11.944728

**Authors:** Wenjun Zhang, Zeliang Chen, Yi Lu, Zhongmin Guo, Yanhong Qi, Guoling Wang, Jiahai Lu

## Abstract

A discrete dynamic model for human epidemics was developed in present study. The model included major parameters as transmission strength and its decline parameters, mean incubation period, hospitalization time, non-hospitalization daily mortality, non-hospitalization daily recovery rate, and hospitalization proportion, etc. Sensitivity analysis of the model indicated the total cumulative cases significantly increased with initial transmission strength, hospitalization time. The total cumulative cases significantly decreased with transmission strength’s decline and hospitalization proportion, and linearly decreased with non-hospitalization daily mortality and non-hospitalization daily recovery rate. In a certain range, the total cumulative cases significantly increased with mean incubation period. Sensitivity analysis demonstrated that dynamic change of transmission strength is one of the most important and controllable factors. In addition, reducing the delay for hospitalization is much effective in weakening disease epidemic. Non-hospitalization recovery rate is of importance for enhancing immunity to recover from the disease.

## 1 Introduction

So far a lot of dynamic models have been developed and used in the mechanic analysis and prediction of animal epidemics. Among them the differential equations based models are the mainstream methods, including SIR model (Kermack and McKendrick, 1927), Anderson-May model (Anderson and May, 1981), the models of Zhang et al. (1997, 2011), etc. Zhang et al. (1997) model was a group of differential-integral equations mainly treating susceptible and infected insect populations. The improved model (Zhang et al., 2011) was composed of nearly twenty differential equations supplemented by other equations. In these models, both susceptible and infected populations were treated as population density, and susceptible population interacts with infected population mainly through feeding on virus on the leaves spread by insects that died from virus infection. Serving as both explanatory and simulation models, they have demonstrated the better performance. Both SIR model (Kermack and McKendrick, 1927) and Anderson-May model (Anderson and May, 1981) include differential equations (correspondingly, difference equations) for susceptible (*s*) and infected (*i*) populations rather than population density, and the two populations interact with each other through the interaction term, p*s*(*t*)*i*(*t*) (Fuxa and Tanada, 1987). Nevertheless, numerous simulation results showed that both of their models are extremely sensitive to some of the key parameters and initial population sizes, especially the infection coefficient, p. Given true parameters and initial conditions, it was so difficult to synchronously obtain realistic results for population size and key time points (such as the peak time) although both of them are better explanatory models for the epidemic dynamics. Furthermore, the important parameters such as incubation period, hospitalization term, etc., were not included in such explanatory models and most of the other models (Chen et al., 2020a). In present paper we developed a generalized discrete dynamic model for human epidemics, and sensitivity analysis and scenario prediction was made, aiming to provide a novel simulation tool for future uses.

## 2 Methods

### 2.1 Model

Suppose the susceptible population is infinite compared to the infected population, i.e, the susceptible population is approximately a constant. The susceptible population can thus be ignored. In addition, the hospital acts as a “black hole”. The hospital accommodates infected cases, and the later recovered from medical treatment and are released to the susceptible population, or die. According to general rules and human operations for disease epidemics, the generalized discrete dynamic model (delay difference equation) for human epidemics is developed as the following

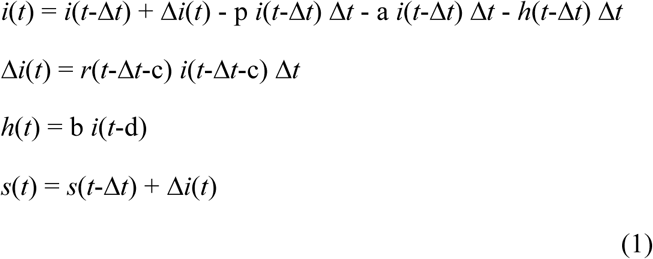

where

*i*(*t*): non-hospitalization cases at time *t*

Δ*t*: time step

Δ*i*(*t*): non-hospitalization new cases at time *t*

*h*(*t*): hospitalization cases at time *t*

*s*(*t*): cumulative cases at time *t*

*r*(*t*): transmission strength of non-hospitalization cases at time *t* in Δ*t*

c: mean incubation period

d: mean time from onset to hospitalization

a: non-hospitalization disease mortality in Δ*t* (0≤a≤1)

b: proportion of non-hospitalization cases for hospitalization (0≤b≤1)

p: non-hospitalization recovery rate in Δ*t* (0≤p≤1)

Equation (1) can be represented by a delay differential equation and an integral

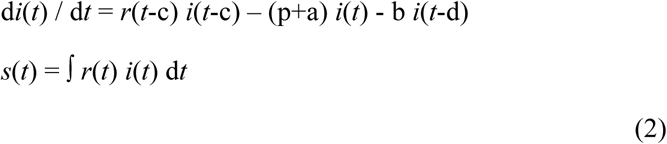

Without losing generality, let Δ*t*=1 (e.g., one day), we have the following dynamic model corresponding to equation (1)

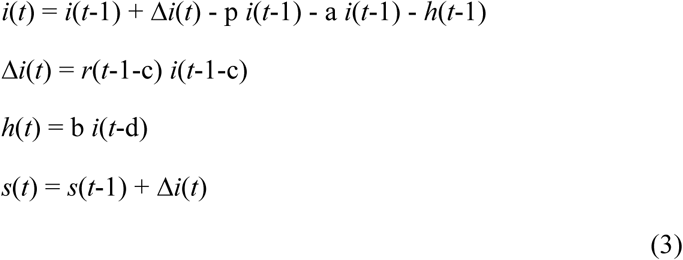

### 2.2 Problems of *r*(*t*), p, a, and b

The transmission strength of non-hospitalization cases, *r*(*t*), is a function of time *t*, dependent upon the type of dynamics of transmission strength of non-hospitalization cases.

As the most occurred type, *r*(*t*) may decline with time for the reasons such as the increase of people’s self-protection, and other transmission-reducing measures adopted by governments and individuals, etc. In this situation, it can be expressed as the linear approximation

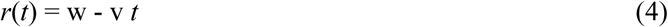

*r*(*t*) can also be represented by other functions. Some of the representative functions include

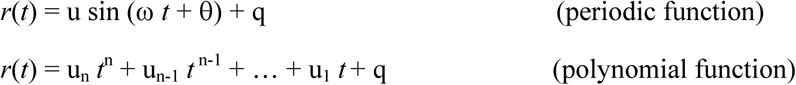

A method to find *r*(*t*) is based on the concept of reproduction number. According to the definition, *r*(*t*) = *R*(*t*) - 1, where *R*(*t*) is the reproduction number at time *t*. if *r*(*t*) = r_0_, then r_0_ = R_0_-1, where R_0_ is the basic reproduction number. In the equation (4), we can take w = R_0_ - 1. Ignoring the parameters b, a, and p, the epidemics will limitlessly increase if R_0_>1, will maintain at a certain level if R_0_=1, and will gradually disappear if R_0_<1.

In addition, we can set the parameters, p, a, and b in equations (1) ∼ (3) as dynamic variables, i.e., *p*(*t*), *a*(*t*), and *b*(*t*) (It should be noted that epidemic dynamics (1) ∼ (3) is a natural epidemics without artificial interference if b=0 or *b*(*t*)=0).

Dynamics of non-hospitalization new cases will be diverse, which is dependent upon the specific functions, *r*(*t*), *p*(*t*), *a*(*t*), and *b*(*t*). For example, the periodic oscillation may occur in certain conditions, etc. A typical dynamic type is illustrated in Fig. 1.

**Fig. 1.**
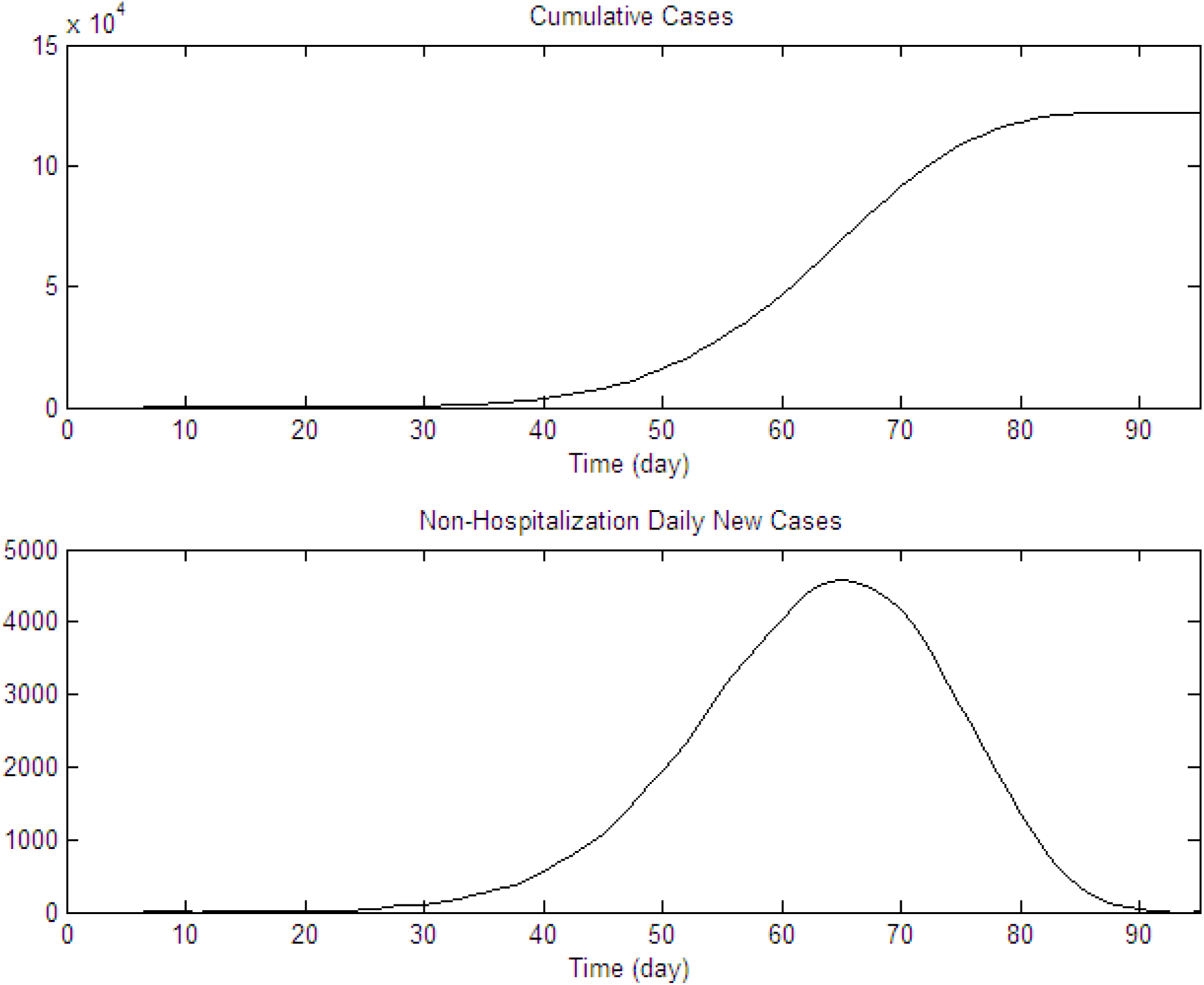
A type of epidemic dynamics produced by equation (3) (based on equation (4)).

## 3 Sensitivity analysis

As the most occurred type, based on equation (4) and some parameters of Novel Coronavirus Pneumonia disease (Chen et al., 2020; Guan et al., 2020; Guo et al., 2020; Liu et al., 2020; Riou and Althaus, 2020), we assume a set of background parametrical values of (c,d,p,a,b,w,v) as (4,3,0.01,0.01,0.9,2,0.019) for sensitivity analysis. The first infection case (*h*(*t*)=1 in equation (3)) was diagnosed on Dec 8, 2019. Considering the mean hospitalization time, d=3, the initial infected population should be 1 on Dec 5, 2019, i.e., *i*(0)=1 (*t*=0: Dec 5, 2019).

### 3.1 Transmission strength

#### (1) Effect of the change of initial transmission strength (w)

Total cumulative cases (TCC) increases exponentially with initial transmission strength (w) (Fig. 2). The 0.1 of increase in initial transmission strength (w) will result in an increase of TCC by 290%, while the 1517% increase of TCC is expected by an increase of 0.2 of w.

**Fig. 2.**
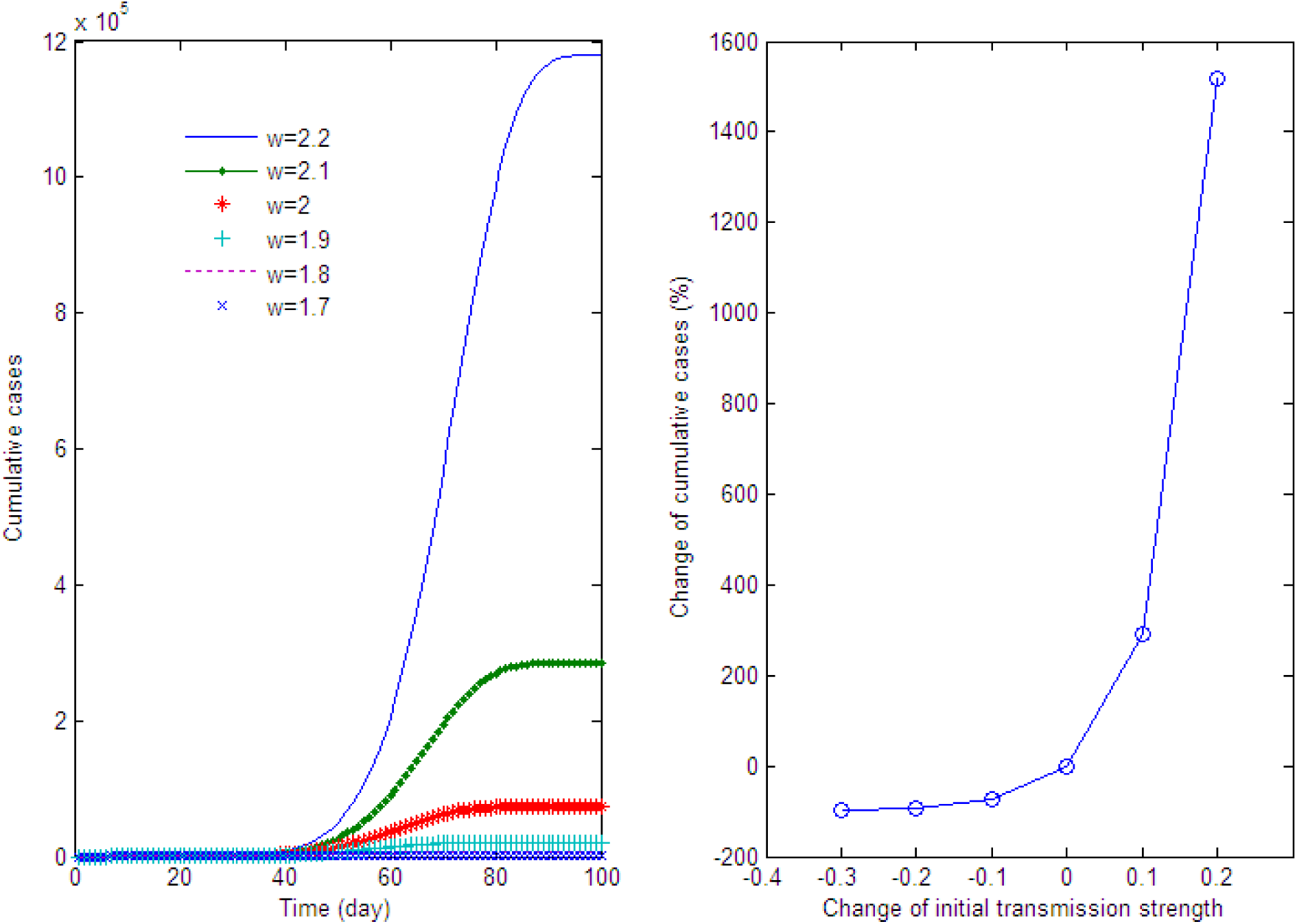
Effect of the change of initial transmission strength (w).

#### (2) Effect of the change of transmission strength’s decline (v)

TCC decreases exponentially with transmission strength’s decline (v) (Fig. 3). The 0.001 of increase in v will result in a decline in TCC by 37%, while the 0.002 of increase in v will result in a decline in TCC by 59%.

**Fig. 3.**
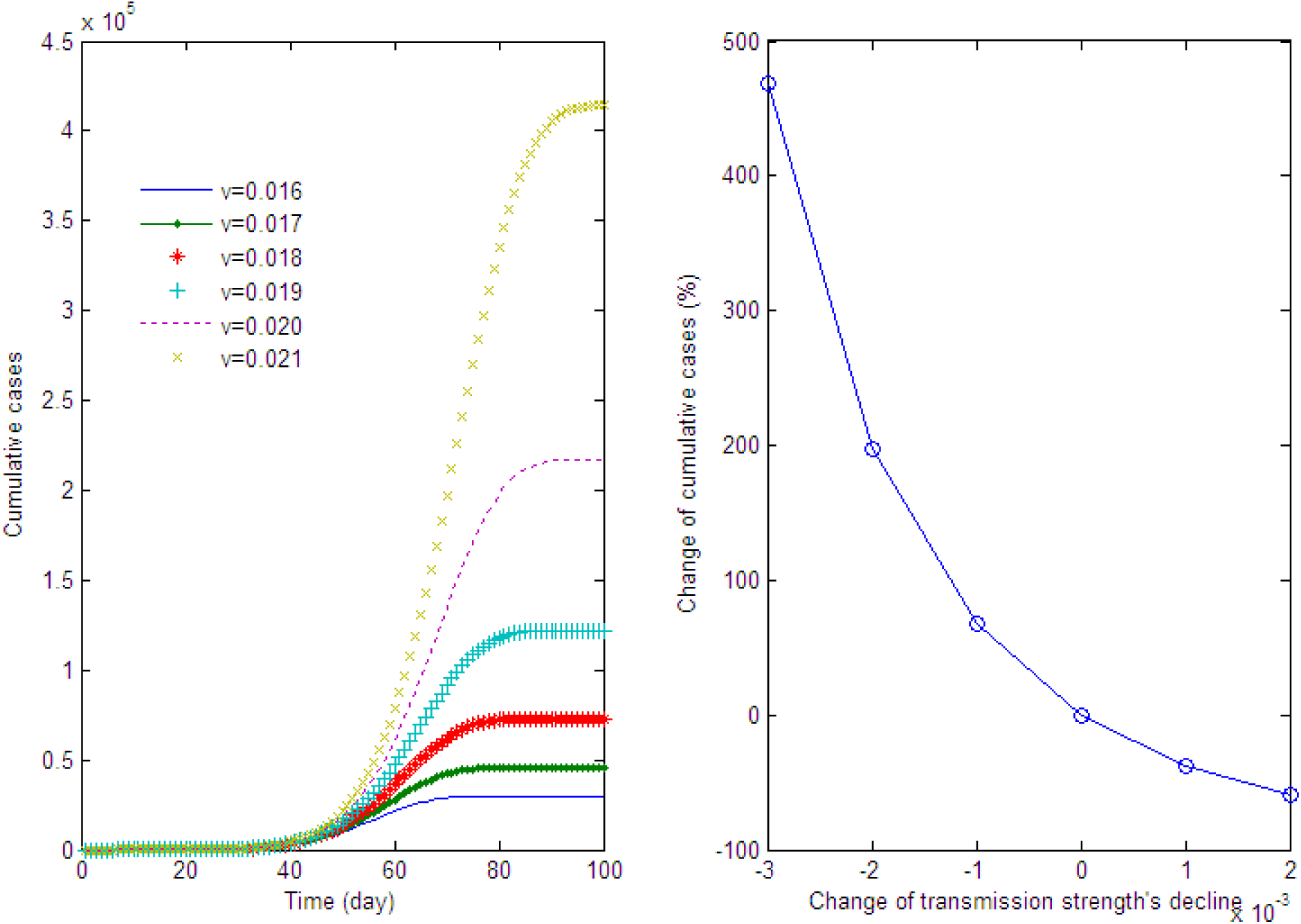
Effect of the change of transmission strength’ decline (v).

### 4.2 Effect of the change of mean incubation period (c)

TCC decreases dramatically with mean incubation period (c) when c is less than a certain value while it slowly increases with mean incubation period (c) if c is greater than the value (Fig. 4). One day’s decrease in mean incubation period (c) will lead to a decline in TCC by 1817%.

**Fig. 4.**
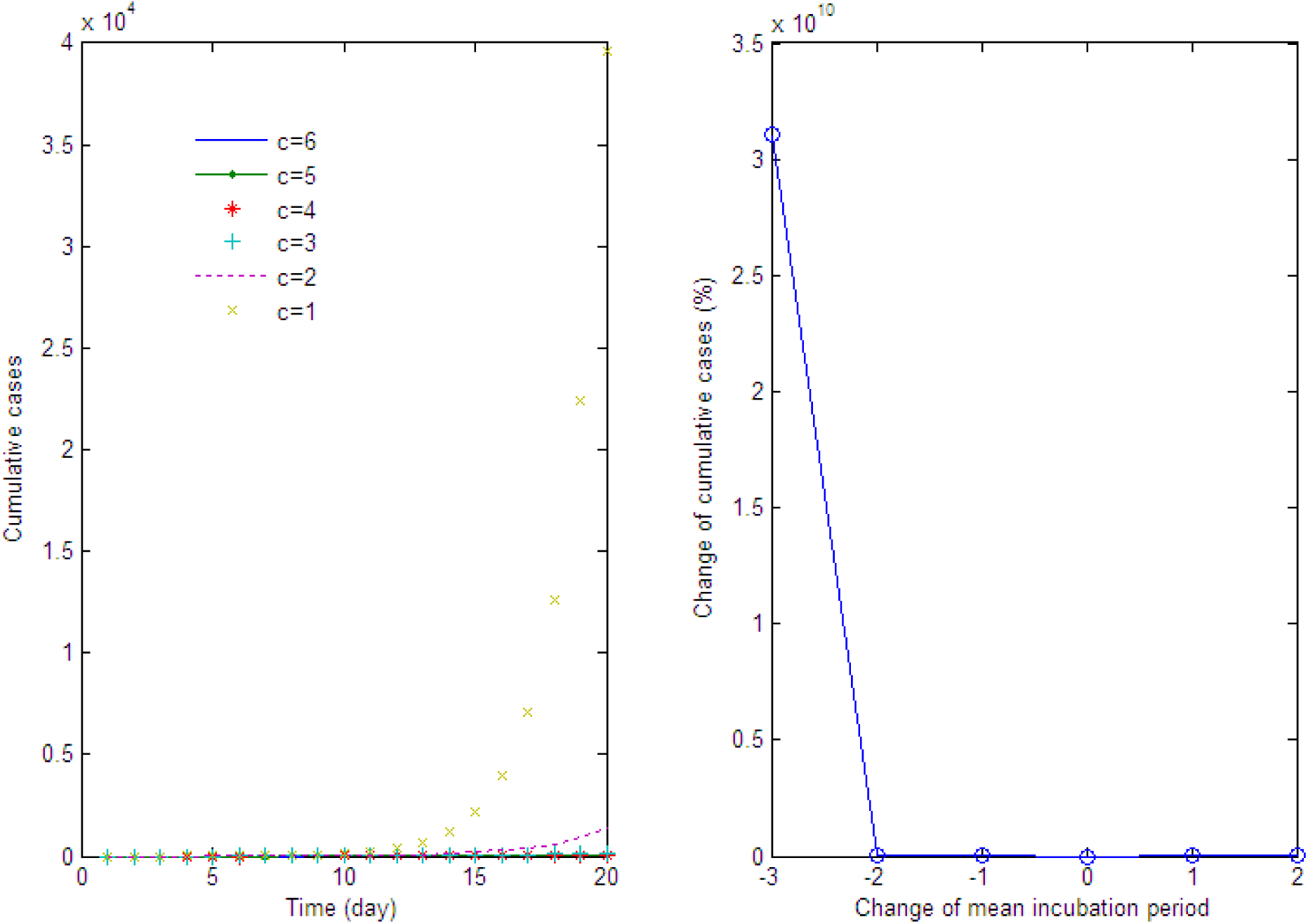
Effect of the change of mean incubation period (c).

### 4.3 Effect of the change of hospitalization time (d)

TCC increases exponentially with hospitalization time (d) (Fig. 5). However, at least 40 to 50% decrease in TCC will appear if hospitalization time is less than 4 days.

**Fig. 5.**
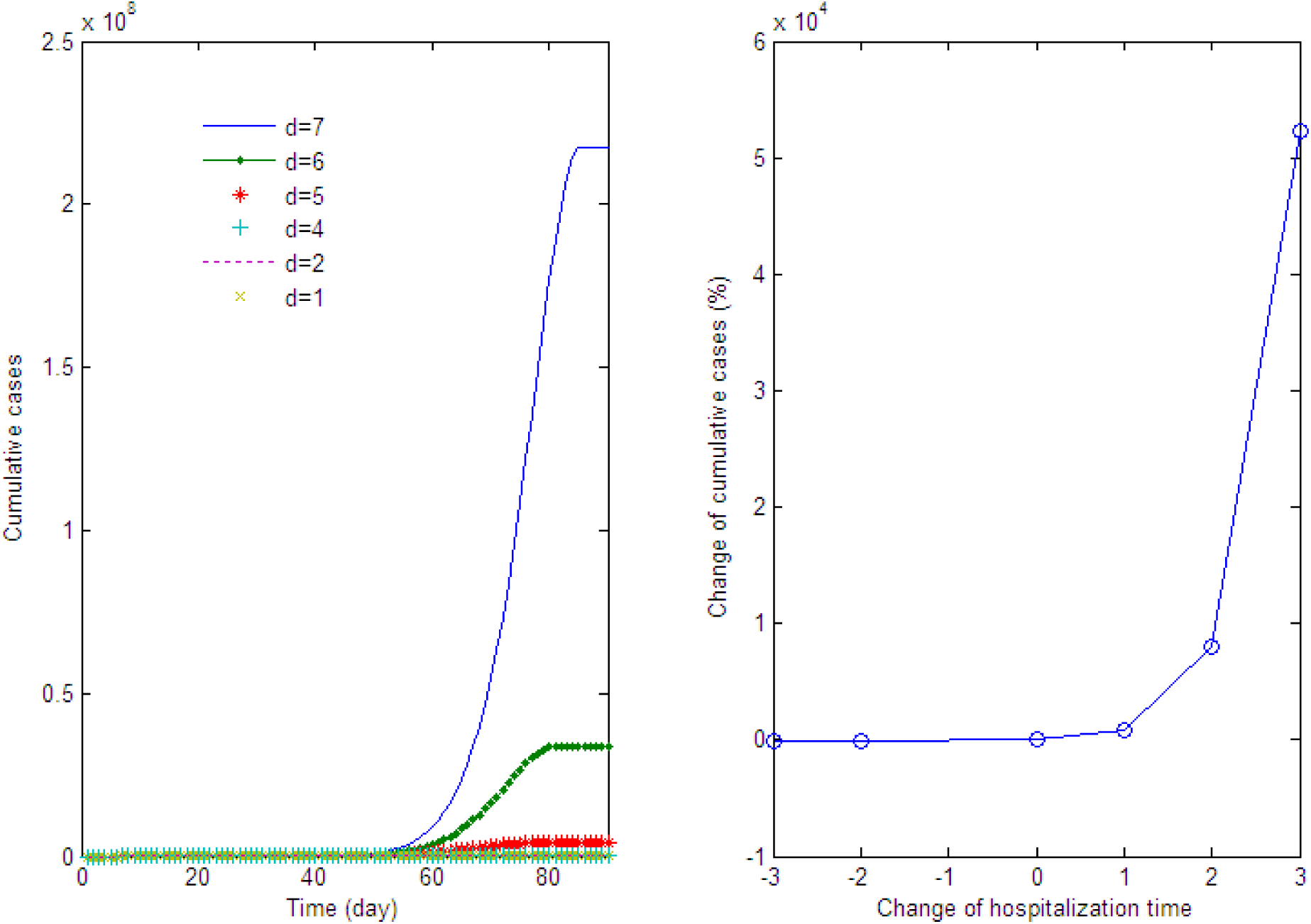
Effect of the change of hospitalization time (d).

### 4.4 Effect of the change of non-hospitalization daily mortality (a) and daily recovery rate (p)

TCC decreases linearly with non-hospitalization daily mortality (a) and non-hospitalization daily recovery rate (p) (Fig. 6 and 7). The 0.005 of increase in p or a will lead to a decline of TCC by nearly 10%.

**Fig. 6.**
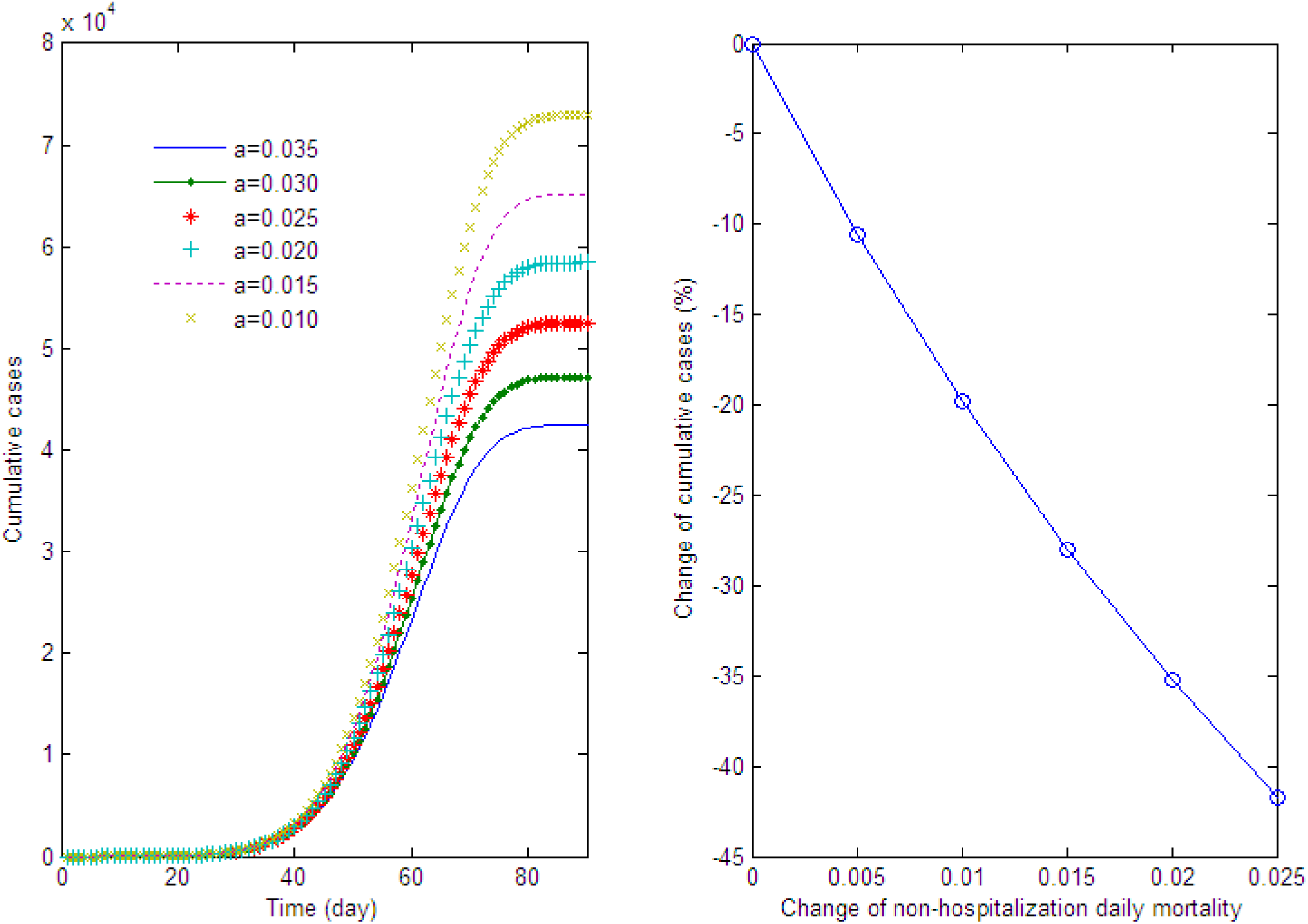
Effect of the change of non-hospitalization daily mortality (a).

**Fig. 7.**
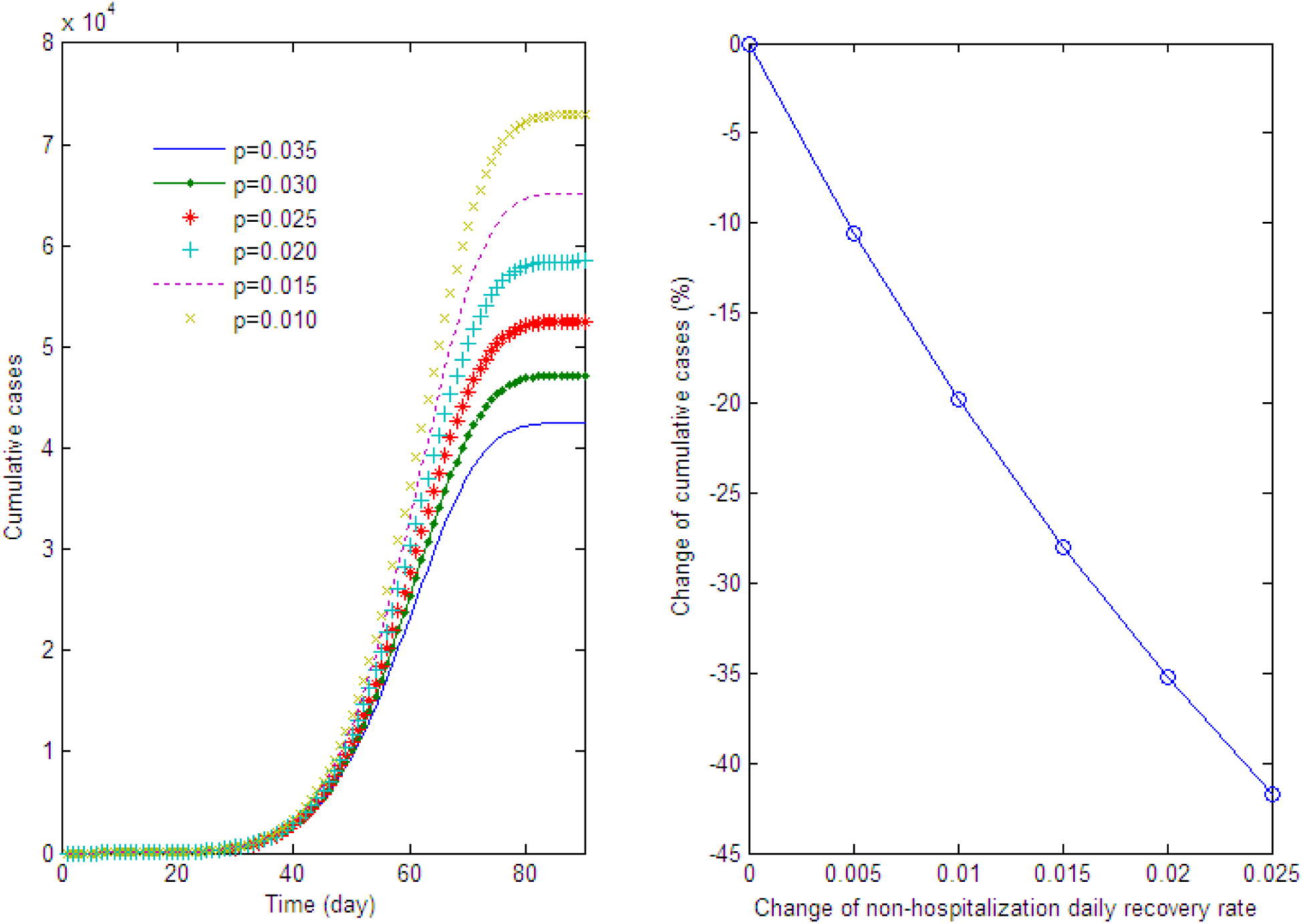
Effect of the change of non-hospitalization daily recovery rate (p).

### 4.5 Effect of the change of hospitalization proportion (b)

TCC decreases exponentially with hospitalization proportion (b) (Fig. 8). The 10% of decrease in hospitalization proportion will lead to the increase in TCC by 318%, and 20% of decrease in hospitalization proportion will lead to the increase in TCC by 1819%,

**Fig. 8.**
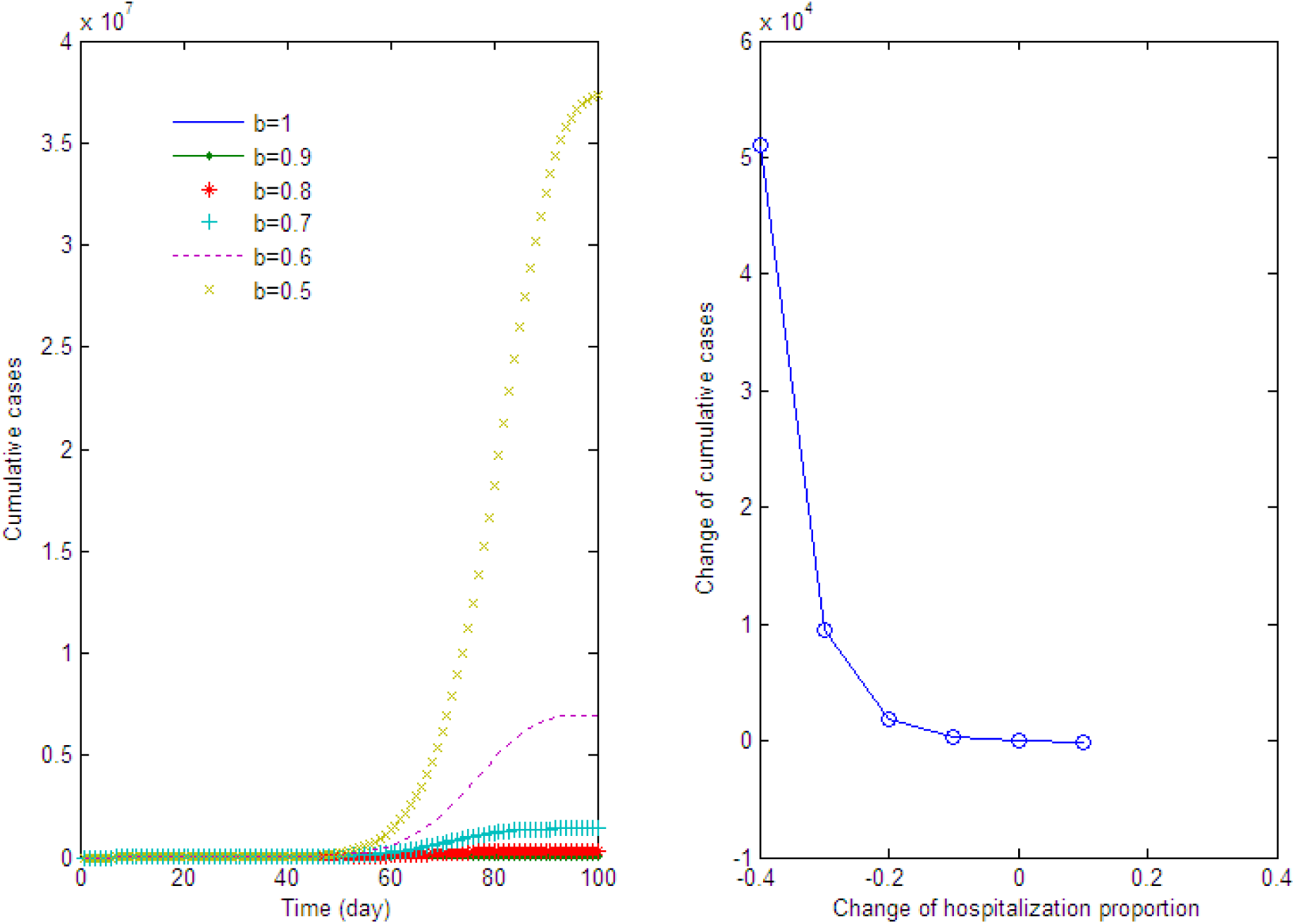
Effect of the change of hospitalization proportion (b).

## 5 Discussion

Sensitivity analysis indicated that almost all parameters have significant influence on the dynamics and final outcome of human epidemics. Among them, the dynamic change of transmission strength (equivalently, the dynamic change of R_0_) is one of the most important and controllable factor. The human epidemics can be deterministically weakened by reducing transmission strength. In addition, the results demonstrated that reducing the delay for hospitalization is much effective in weakening disease epidemic. Sensitivity analysis on non-hospitalization recovery rate reminds us the importance for enhancing immunity to recover from the disease. Mean incubation period is another key factor in determining disease epidemics. In a certain range, a short incubation period may lead to the rapid development and serious outcome of epidemic disease.

As shown in sensitivity analysis, the model performance depends upon the exact parametrical values. How to obtain the exact parametrical values for the disease is the basis for the better simulation and prediction of model performance. On the other hand, some parameter, e.g., incubation period falls in an interval rather than a deterministic value. Therefore, a comprehensive analysis based on scenario prediction is necessary to obtain the most reliable prediction.

In present model, both asymptomatic infection and nosocomial infection were ignored due to their general insignificance in most epidemic diseases. For some diseases, however, these infections would play an important role and should be included in the model. Further, more complex models that include other factors or processes, e.g., network properties, can be developed for specific uses (Shams and Khansari, 2019; Chen et al., 2020a).

